# Capuchin monkey rituals: an interdisciplinary study of form and function

**DOI:** 10.1101/2020.02.21.958223

**Authors:** S. Perry, M. Smolla

## Abstract

Many white-faced capuchin monkey dyads in Lomas Barbudal, Costa Rica, practice idiosyncratic interaction sequences that are not part of the species-typical behavioural repertoire. These interactions often include uncomfortable or risky elements. These interactions exhibit the following characteristics commonly featured in definitions of rituals in humans: (1) they involve an unusual intensity of focus on the partner, (2) the behaviours have no immediate utilitarian purpose, (3) they sometimes involve “sacred objects”, (4) the distribution of these behaviours suggests that they are invented and spread via social learning, and (5) many behaviours in these rituals are repurposed from other behavioural domains (e.g. extractive foraging). However, in contrast to some definitions of ritual, capuchin rituals are not overly rigid in their form, nor do the sequences have specific opening and closing actions. In our 9,260 hours of observation, ritual performance rate was uncorrelated with amount of time dyads spent in proximity but is (modestly) associated with higher relationship quality and rate of coalition formation across dyads. Our results suggest that capuchin rituals serve a bond-testing rather than a bond-strengthening function. Ritual interactions are exclusively dyadic, and between-dyad consistency in form is low, casting doubt on the alternative hypothesis that they enhance group-wide solidarity.

## 1 INTRODUCTION

A long-term field study of white-faced capuchin monkeys in Lomas Barbudal, Costa Rica has yielded a rich observational record of highly idiosyncratic interaction sequences not found in the species-typical behavioural repertoire. Here, we (a) give an account of these puzzling, apparently non-utilitarian social interaction sequences practiced by some, but not all, capuchin monkey dyads, (b) determine whether these behaviours qualify as rituals according to definitions of ritual in various disciplines, and (c) test hypotheses regarding the possible function or communicative role these interactions might serve, by examining the qualities of the behaviours themselves and the characteristics of the dyads performing them.

We define capuchin rituals as “learned behavioural sequences with no obvious immediate utilitarian purpose, composed of behavioural elements repurposed from other parts of the behavioural repertoire, characterized by a high degree of attentional focus by one or both partners on the other’s body and/or a (“sacred”) object jointly handled by the interactants.” However, we also consider whether these interaction sequences we define as rituals contain elements from other researchers’ definitions of ritual (see Table 1).

**Table 1:** Elements commonly found in definitions of ritual across disciplines.

| Definitional feature of ritual | Intellectual tradition(s)* | Reference examples | Proposed ritual function** | Present in capuchins |
| --- | --- | --- | --- | --- |
| Distinction between the sacred and the profane | ANTH, SOC | Bell, 1997; Durkheim, 1912; Goffman, 1967; Rappaport, 1999 | b, c, e | ? |
| Shared group emotion | SOC | Collins, 2004; Durkheim, 1912 | e | No? |
| Feedback between mood and joint focus; shift in emotional energy within a dyad | ETHO, SOC | Collins, 2004; Zahavi, 1977 | c, d, e | Yes |
| Breaking ritual proprieties results in moral uneasiness | SOC | Goffman, 1967; Rappaport, 1999 | b | Probably not |
| Repurposing of behaviors evolved for other purposes | ETHO | Huxley, 1914; Smith, 1977 | c | Yes |
| Exaggeration and stereotypy (comparable to formalism) | ETHO, PSY, ANTH? | Bell, 1997; Boyer and Liénard, 2006a; Merker, 2009; Rappaport, 1999 | a, b, c | Only slightly |
| Traditionalism, i.e. repetition of activities from an earlier period | ANTH | Bell, 1997 | b, c | Yes |
| Disciplined, rigid repetition, often rhythmic | PSY | Bell, 1997; Boyer and Liénard, 2006a; Merker, 2009; Smith, 1977 | a, b? | Only slightly |
| Lack of rational motivation, no obvious immediate function | ANTH, ETHO, PSY, SOC | Boyer and Liénard, 2006a; Rappaport, 1999 | a, b, c, d, e | Yes |
| Compulsiveness | PSY | Boyer and Liénard, 2006a | a | No? |
| Framing of the act | ANTH | Bell, 1997; Ortner, 1973 | (various) | No |
| Symbolism | ANTH, SOC | Bell, 1997; Collins, 2004; Durkheim, 1912; Goffman, 1967; Ortner, 1973; Rappaport, 1999 | b, c, e | Probably only during the ritual |

\* ANTH = anthropology, SOC = sociology, ETHO = ethology, PSY = psychology
\*\* a = detection of and reaction to inferred fitness threats,
b = transmission and maintenance of social norms,
c = communication about/definition of social roles,
d = testing of dyadic social bonds,
e = establishing society-wide solidarity

### 1.1 Definitions across academic disciplines

There is very little consensus among researchers, either within or between disciplines, about what a ritual is. However, several features appear repeatedly in these definitions. In Table 1, we present a sample of some common definitional features of ritual associated with particular disciplines. The features selected as being important in these definitions probably have some loose association with the putative function of ritual that is attributed by the researchers. The fourth column in Table 1 lists the putative function of ritual that is named in studies for which this feature is an essential part of the definition of ritual. Note, however, that in some definitions, certain features are claimed to be indispensable and others optional, and that the claimed link between definitional feature (form) and function is made explicit in some, but not all of these studies, most of which were not designed with such an evolutionary analysis in mind. This is not an exhaustive literature review of the many elements found in definitions of ritual, nor of the researchers using these definitions. Instead, it is meant as a rough guide demonstrating some of the cross-disciplinary linkages in definitions and theorizing regarding the function of ritual that we came across.

Here we describe how the characteristics of the capuchin monkey behaviours that we designate as “rituals” in the Results section correspond to these proposed attributes in our definition:

1. *Quality of attentional focus:* The behaviours described here are prolonged (dyadic) social activities sometimes lasting up to an hour, involving a high degree of focus by one or both partners on the other’s body and/or actions, and/or an object jointly handled by the interactants, thereby diverting attention away from normal activities such as foraging or vigilance (as in Rossano’s definition of ritual (Rossano, 2011)). This intense focus on a particular partner, to the exclusion of other group members, is reminiscent of Collins’ theory of interaction rituals (Collins, 2004), in which degree of attentional focus and “emotional energy” directed to a partner provides information to the recipient about its current relative value to the actor.
2. *Lack of immediate utilitarian purpose:* These behaviours have no obvious utilitarian purpose, i.e. they do not seem to enhance food-acquisition, safety or health in any obvious, immediate way, though they may serve a communicative purpose.
3. *Sacred objects:* Similarly, the objects (and partner body parts) handled in some of these rituals have no utilitarian value, e.g. they are neither food nor tools. Whether or not they have any symbolic value, qualifying as “sacred objects,” depends on how this term is defined (see discussions by Durkheim (1912), Goffman (1967) and Collins (2004)), but it seems unlikely that the objects retain this symbolic value after the ritual is over.
4. *Learned behaviours:* We infer that these behaviours have a learned component because they are not performed by all individuals in the population, and are not produced during early development. They could hypothetically appear in an individual repertoire via innovation or social learning (probably via ontogenetic ritualization, Tomasello et al. 1989).
5. *Repurposing of behavioural elements found elsewhere in the repertoire:* Scholars of both ritual and play (Goffman, 1967; Smith, 1977; Watanabe and Smuts, 1999) have noted that complex animal rituals, like human rituals, often involve the transfer of behavioural elements and stimuli from one behavioural domain to a new (social) context.

Importantly, we do *not* emphasize rigidity of form or repetition in our definition, although this attribute comprises a core feature of many scholars’ definitions of ritual (e.g. Bell 1997; Boyer and Liénard 2006b), nor is function a critical part of our definition. However, we did choose to focus on those definitional aspects used by scholars who study the implications of ritual for social relationships (Table 1). Although some degree of repetition and constancy of form is clearly involved in ritual, both human and nonhuman, we argue that rituals may serve multiple functions, and that the optimal degree of flexibility vs. rigidity in form may depend on the function of the ritual.

There was insufficient space to discuss definitional aspects needed for testing some alternative hypotheses, e.g. that the capuchin rituals we describe function to detect and react to inferred threats (Boyer and Liénard 2006), or the idea that capuchin rituals contain symbolic content relevant to detecting or enforcing social norms violations. However, the lack of precisely repeated, rhythmic, compulsive actions is incompatible with the Hazards Precautions Hypothesis and the social norms hypothesis. Furthermore, there is no obvious display of moral outrage or shame in response to deviations from the typical form that a dyad’s ritual has, nor is there currently any solid evidence for social norms in capuchins.

### 1.2 Hypotheses to be tested

Researchers of nonhuman primates have developed several hypotheses regarding the relationship between social relationships and rates of greeting rituals. Some hypotheses state that ritual performance is necessary to establish or maintain social bonds, perhaps by defining social roles and negotiating the terms of a social relationship, and predict that ritual performance will be associated with higher frequencies of time spent together, affiliative behaviours, or cooperation (De Marco et al., 2014). Others state that these rituals are a way of testing important relationships critical to enhancing fitness (De Marco et al., 2014; Perry et al., 2003; Watanabe and Smuts, 1999; Whitham and Maestripieri, 2003; Zahavi, 1977), which leads to the predictions that bond tests will be more frequent when (a) there is a dearth of information regarding the state of the relationship, (b) there is good reason to believe that the relationship is undergoing change (e.g. during a rank reversal), and (c) the bond is solid enough that a Zahavian bond test won’t be extremely risky, yet not so secure that there is no need to test it at all.

To gain insight into the function of these rituals, we test the following hypotheses and predictions

1. Rituals serve to establish and maintain social bonds:
  a. Rituals will be more frequent in dyads that (i) spend the most time in proximity, (ii) have higher relationship quality, and (iii) cooperate most often in coalitionary aggression.
2. Rituals serve as Zahavian tests of social bonds (Zahavi, 1977):
  a. Behavioural elements will entail some risk and/or discomfort.
  b. Rituals will be more frequent when state of a relationship is unclear (i.e. there should not be a positive linear correlation between rate of ritual performance and time spent together, but rather, higher rates at intermediate amounts of time spent in association).
  c. Rituals are predicted to be most often performed in dyads with good enough relationships to feel comfortable performing the intimate ritual, but not in relationships so completely free of conflict that they require no testing (i.e. highest rates of ritual performance at upper intermediate values of relationship quality rather than at the highest end of the distribution).
3. Participation in rituals promotes group-wide solidarity:
  a. Rituals are expected to be performed simultaneously by many monkeys at once, exhibiting a form that is consistent among group-mates

## 2 STUDY SPECIES AND METHODS

### 2.1 Study population

Our subjects are wild, well-habituated white-faced capuchin monkeys (*Cebus capucinus*), residing in and near Lomas Barbudal Biological Reserve, in the tropical dry forests of northwestern Costa Rica. This population has been studied since 1990 by S. Perry and collaborators (see (Perry, 2012) for more details on the natural history of this species, and Perry et al. 2012 and Perry and Manson 2008 for information on this longitudinal project, including the methods). White-faced capuchins are extraordinarily large-brained, long-lived New World primates living in stable multi-male, multi-female groups, characterized by female philopatry and male parallel dispersal; i.e. both sexes can maintain long-term bonds with same-sexed kin (Perry, 2012). Their social behaviour is complex and characterized by a rich repertoire of signals for communicating about their social relationships, including both species-typical vocalizations and gestures, and innovative/learned gestures (Perry, 2012). Cooperative interactions and alliances are key to the reproductive success of both sexes and pervade many aspects of capuchins’ lives (Perry, 2012).

### 2.2 Data collection

Observers were instructed to record, in minute detail, descriptions of any social interaction (during focal and *ad libitum* observation) that was not composed exclusively of standard (i.e. species-typical) items in the ethogram in their normal context. Interaction descriptions were recorded in the field and later transcribed into a daily spreadsheet. Whenever possible, interactions were videotaped. Observers recorded participants’ posture/bodily orientations, gaze directions, which body parts were in contact, any physical object that was handled as part of the interaction, and the social context (e.g. whether other monkeys were in proximity and whether they were paying attention). During *ad lib* observations, interaction start times were sometimes missed. Descriptions varied somewhat in level of detail, as the unpredictable form of these innovative interactions made it difficult to devise appropriate inter-observer reliability measures. To increase reliability, two observers typically collect data from two different locations, thus mitigating the problem of foliage obscuring some parts of the interaction.

### 2.3 Data set

The new data presented here are from “Flakes” (FL) group, which fissioned from Abby’s group (the original study group) in late 2003. Here, we analyse 9,260 observation hours of data collected between February 1, 2004, when the group had become demographically stable, until October 11, 2018. FL group was composed of two matrilines, headed by matriarchs who are probably cousins, and contained five immigrant males, who arrived singly at different times during 2003-2004; two of these shared a natal group, and three were from outside the study area. These immigrant males seemed to be 8-12 years old at the start of 2004. Over the course of the 15 years of observation, Flakes group included 53 individuals, ranging from 9-30 members at any given time (six monkeys were excluded from the analysis who died prior to 6 months of age). The data set consists of 446 social interaction “rituals” and 6 failed attempts of monkeys to elicit joint interaction in a ritual. Thirty-seven (79%) of the 47 group members included in the analysis (17 of 20 females and 20 of 27 males), and 17% of the 762 co-resident dyads (40 female-female, 47 male-female and 46 male-male dyads) engaged in at least 1 ritual. Only 19 individuals (6 female, 13 males) comprising 32 dyads (25 male-male and 7 male-female) participated in the most complex rituals (i.e. “games”). As more peripheral monkeys are more often missed in group scans, and because only a few individuals were the subjects of focal observations, total observation time varies among individuals. Observers spent much of each day collecting focal follows, so the probability of detecting rituals performed by focal subjects was higher than for non-focal animals. To correct for observation effort, we summed the number of group scans and point samples (collected at 2.5-min intervals during focal follows) for each member of the dyad on days when both members of the dyad were co-resident in FL group.

### 2.4 Measures

To test hypotheses 1 and 2 we require measures for: physical proximity, relationship quality, and coalition formation. Our measure of physical proximity is based on “group scans” in which researchers wandered through the group, recording distance between the scanned monkey and other monkeys in proximity to it. We scored two individuals as being in physical proximity if they were within 40cm of one another (equivalent to an adult male body length, from nose to tail base).

Our measure of relationship quality (RQ) is based on observed social interactions during focal follows (when available) and *ad libitum* observations. The standard social interaction repertoire included 79 behaviours (some dyadic and some triadic) with clear emotional valences, i.e. participation would elicit at least in one of the participants positive or negative emotions, thereby affecting the “emotional energy” (sensu Collins (2004)) or the emotionally-mediated “book-keeping” of the rates and qualities of interactions (Schaffner and Aureli, 2002) expected to influence the quality of future interactions of that dyad. Grooming, playing, and forming a coalition are among the 42 behaviours expected to have a positive impact, whereas aggression and submission are among the 37 behaviours expected to have negative impact on relationship quality. We determined the relationship quality index (RQI) in the following way: we aggregated data into 10-minute chunks. For each dyad and for each chunk, we assigned a score of 1 if there was at least one positive behaviour, and 0 if not. We repeated this for negative behaviours. The RQI for a given dyad-year consists of (a) the number of time-chunks with one or more positive impact behaviours, divided by the sum of (a) plus (b) the number of time-chunks with one or more negative impact behaviours. Thus, RQI = 0 represents exclusively negative and RQI = 1 exclusively positive “emotional energy”.

Our measure of coalition formation is based on *ad libitum* data, using only incidents of “overlords”, “cheek-to-cheek” postures, and “embraces” (defined in Perry 2012) against a common conspecific opponent. Although coalitions are fairly conspicuous behaviours, there is nonetheless some tendency to underreport coalitions from peripheral group members. We accounted for this by creating an offset variable consisting of the number of group scans collected for individual A and individual B on days when they were co-resident. Because *ad libitum* observations were collected primarily as observers were wandering through the group collecting group scans, this should be a fairly accurate representation of the observability of these individuals. Given the sometimes large number of group scans per individual, we divided the total group scans by 1000, and so, our coalition index represents the number of observed coalitions per 1000 observations. The reason for lumping behaviours within 10-minute time intervals is that individuals likely adjust emotional energy in a more bout-like fashion. E.g. they recall losing a fight to a particular monkey, but probably they don’t feel much worse for having been slapped 7 times, as opposed to 4 times, within the time period of a 4-minute fight.

### 2.5 Social networks

To analyse how social network position affects ritual performance, and how this relates to the network of ritual participants, we created three networks, based on proximity, rituals, and games (Figure 1a-c respectively). For the social network, we calculated the proximity index, PI, as:

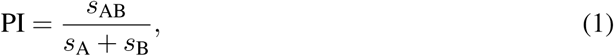

**Figure 1:**
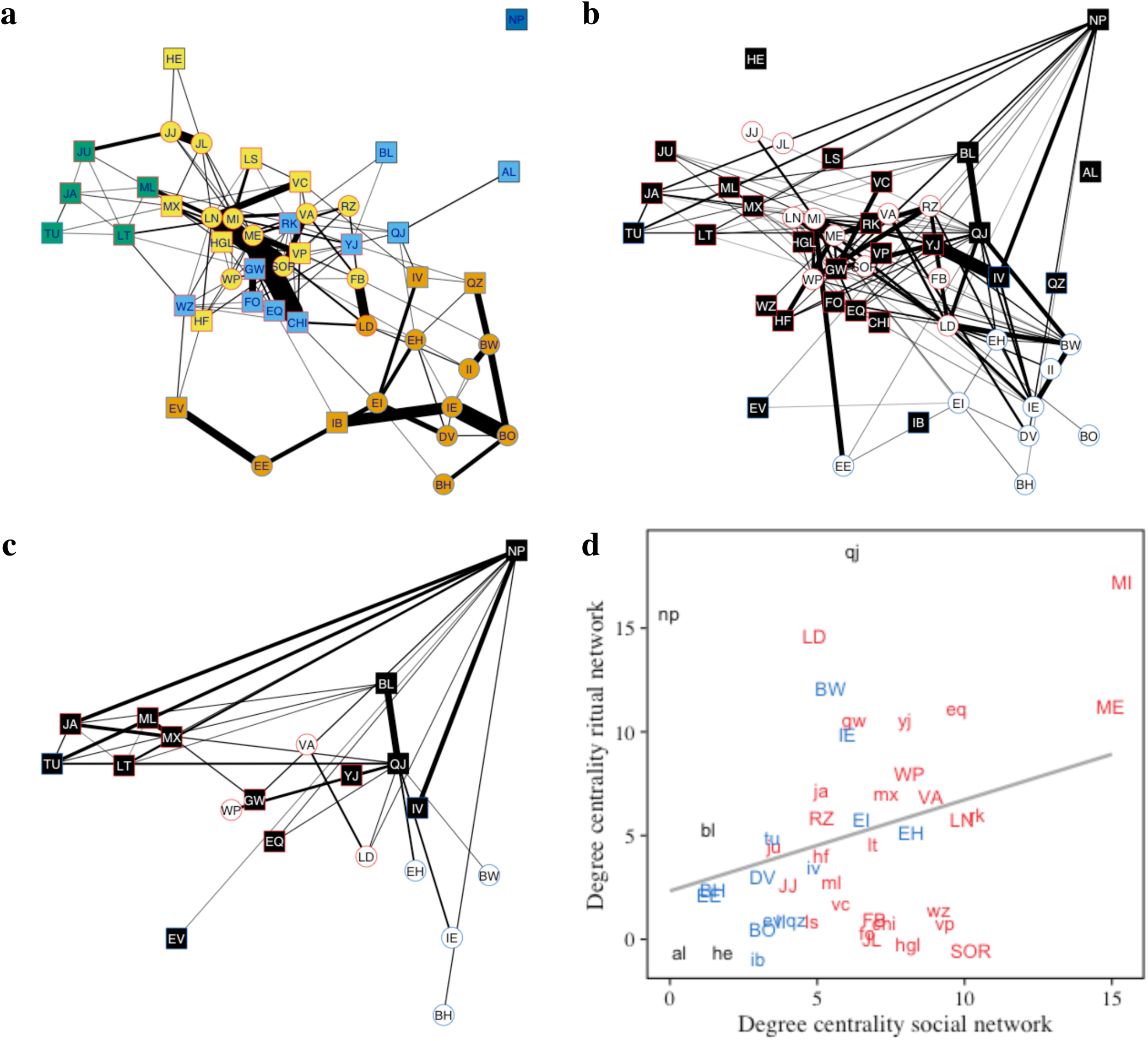
A comparison of the capuchin social network structure and the ritual and game interaction network. The social network of Flakes group (a) shows a slight separation into the two matrilines (red and blue vertex boundaries). The vertex colours indicate to which cluster the individual belongs, based on a greedy community detection algorithm. In all network plots, the vertex shape indicates females (circle) and males (squares), and edge width indicates the strength of interaction or proximity. Inspecting the frequency of rituals among dyads, we find that strong proximity relations (a) do not necessarily indicate high ritual performance frequency (b). This is also true for the games network (c). When we correlate the degree centrality in the social network with the degree centrality in the rituals network (d), we find that the number of individuals with whom a monkey performs rituals (y-axis) is uncorrelated with the number of individuals they regularly interact with (degree centrality, x-axis). There is a slight upwards trend, but it is not significant (regression line in grey, adjusted R^2^ = 0.06, p = 0.06). Males are represented by lowercase letters and females by uppercase letters. Members of the two matrilines are represented by blue (subordinate) and red (dominant), and immigrants by black letters. Note that for ease of comparison, node positions (based on the Fruchterman-Reingold layout) were conserved in all network plots (a-c).

where *s*_AB_ is the number of scans where monkey A and monkey B were ≤1 body length away from each other, and *s*_A_ and *s*_B_ is the number of group scans where each monkey was spotted (together or separate). We only consider scans where both individuals were *co-resident*, that is, they resided in the same group at this day. This avoids inflating the denominator with scans where the monkeys resided in completely different groups, which can be potentially far apart. For the ritual network, we used the number of times a dyad has been found to perform rituals relative to the number of times they have been observed together. We did the same thing for games network.

We used these weights (where 0 means a dyad was never seen together or has never performed a ritual together, and 1 means a dyads has always been spotted together or have performed rituals every time they were observed) to create an adjacency matrix, which lists these weights for every possible dyad. We then created the networks based on these matrices. We used R (Team, 2019), and the igraph package (Csardi and Nepusz, 2006) to create and analyse the networks.

### 2.6 Statistical analysis

This data set was challenging to analyse for five reasons: (1) Most dyads had values of zero, for ritual counts and counts of coalition formation; (2) Dyads are not independent of one another because the same individual can be a member of multiple dyads; (3) There are missing values when constructing a matrix in which each monkey’s name is represented in the row and column headings, because not all possible dyads were co-resident in the group; (4) The use of *ad libitum* data for calculating ritual and coalition rates was a problem because we do not have precise estimates of how much time each individual was observed, and not all individuals are equally easy to observe; and (5) Behavioural sampling density was insufficiently high to permit subdivision of the data set into chunks of time that we intuitively thought would be appropriate for tracking temporal changes in relationship quality and ritual performance rate. We did not find a single analytical approach that satisfactorily dealt with all of these issues. Therefore, we analysed the data set in several different ways. See the Appendix section A3 for additional information about the alternative analytical approaches we tried before settling on MLPE (Maximum Likelihood Population Effects) models as our primary statistical approach.

The data were analysed in a series of three MLPE models, a class of linear mixed effects models that originated in landscape genetics. Like a Mantel test, MLPE models (Clarke et al., 2002) assess the relationship between two matrices. However, the mixed effects parameterization (specifying the covariance structure of the matrices) accounts for non-independence among pairwise data in each matrix (Shirk et al., 2018). Additionally, we could remove dyads that were non-co-resident, which avoided the use of structural zeros. The independent variable was the proximity index, the relationship quality index (RQI), or coalition count, divided by the sum total of group scans of the two coalition partners (as an observability adjustment). In all cases, the outcome variable was the count of rituals performed by this dyad, with the exposure being the sum total of group scans and point samples collected on the two partners in the ritual. Sample sizes of dyads were slightly smaller for the RQI model as a few dyads did not interact. We dropped infants under 6 months of age from the coalition model, since they were never old enough to form coalitions. We use the MLPE rga() function of the ResistanceGA package in R (Peterman, 2018).

## 3 RESULTS

### 3.1 Description of the behavioural phenomenon

Most of the social interactions that comprise a dyadic relationship in white-faced capuchins consist of species-typical interactions common to primates generally: e.g. grooming, hugging, and rough-and-tumble play (chasing, wrestling, biting, hitting, “play face”), submission (cowering, avoiding), aggression, infant care behaviours and sexual interactions, plus a few species-specific behaviours such as coalitionary recruitment signals, courtship “dances” and vocal signals of benign intent or aggressive intent (vocal threats)(Gros-Louis et al., 2008). However, in addition to these species-typical behaviours, white-faced capuchins often invent new forms of social interaction, devising rituals that are often unique in their subtle details to a specific individual or dyad (Perry et al., 2003, 2017). There is inter-individual variation in the propensity to invent such rituals; in a prior 5-year study of innovation in this population, only 84 of 234 individuals (36%) were members of dyads that invented a new social interaction ritual (Perry et al., 2017).

The following behavioural elements were commonly included in novel social rituals created by the monkeys in Flakes group:

1. Inserting a finger into the orifice of a social partner (e.g. mouth, eye, nostril, or ear), or vice versa (inserting the partner’s digits into one’s own orifices),
2. Prying open a mouth or hand to conduct a detailed inspection of its contents,
3. Passing an object (e.g. bark, leaves, flower, stick, green fruit, or hair plucked from the partner’s body) back and forth from one partner to another, taking turns at the role of holding the object in hand or mouth, and extracting it (also with hands or mouth), in a very gentle “tug-o-war”,
4. Clasping of hands, often with fingers interlaced,
5. Cupping the hand over some part of the partner’s face,
6. Sucking on some appendage belonging to the partner (e.g. tail, finger, toe, ear, nose, or sometimes a clump of hair),
7. Using the partner’s back or belly as a drum to create loud, rhythmic noises.

Note that elements 1-3 above seem to be borrowed from the extractive foraging repertoire and applied to a partner’s body rather than to a substrate potentially containing food. The repurposing of elements from one portion of the behavioural repertoire in another section of the repertoire is commonly discussed as a feature of rituals by early researchers of animal ritual (Smith, 1977).

Of the 446 individual instances of rituals described in our sample, 49% involved placing the fingers in or on the nose, 54% involved insertion of fingers in the partner’s mouth, 14% involved passing a “toy” back and forth between mouths or hands, 5% involved biting hair out of a partner who then tried to retrieve it, 7% involved insertion of fingers into a partner’s eye, 7% included “dental exams,” 1% included “back-whacking,” and 4% involved some creative way of kissing, sucking or chewing on a partner. Many rituals included additional features that were more idiosyncratic to an individual or a dyad.

The most complex interaction sequences were the “games” that involved extracting an object from the hand or mouth of the partner (see Appendix section A1.1 for a video clip and a transcription of the interaction sequence). A particularly striking feature was the focus on physical objects (“toys”) that were extracted from interaction partners’ bodies. Sometimes partner 1 would bite tufts of hair out of partner 2, who would then pry open the mouth of partner 1 to recover the hair. Using motor patterns typical of extractive foraging, the hair would then be passed back and forth amicably between the two partners. Other times, non-edible portions of plants were used as the game objects. Note that these objects had no nutritional value, and the monkeys were surrounded by similar objects, which could be more readily obtained. But it seemed that the object acquired value by virtue of the fact that monkey 1 had it in its possession (i.e. it acquired “sacred object” status by virtue of the fact that it was being used in this ritual). The two monkeys would focus their attention on this object for several minutes (usually 10-30 min).

These interactions are readily interpretable from Heesen et al.’s (2017) framework that views social play as joint action, i.e. interactional achievements whereby the participants create a sense of togetherness. They describe three phases of these interactions, including formalized openings and closings, which capuchin rituals generally lack. Instead, our monkeys almost always began and ended their interactions by merely approaching and leaving their partners. However, capuchin ritual behaviour typically includes the characteristics of the middle section (“main body”) of Heesen et al.’s sequence, described as negotiation of continuation of the activity, changes in type of interaction, role reversals, suspension of activities, and re-engagement of partner’s attention to the prior activity. Our subjects often initiate role reversals or changes of activity by explicitly moving their partner’s hands to the part of the body where they want them to be. There are frequent examples of re-engagement of the partner’s attention, both in these rituals and in coalition formation (outside the context of rituals). In the toy and hair “games,” one partner will attempt to re-engage the attention of a partner whose attention has wandered, by spitting out the object and explicitly showing the partner that they have it, before either inserting it in their own mouth, or holding it in front of the partner’s mouth. This is usually successful in re-establishing mutual participation. In a coalitionary context, when there is an asymmetry in affect and participation in attacking an opponent, the angrier monkey will sometimes tug on the body parts of the ally or bounce ferociously while in body contact with the ally, presumably to rev up the partner’s enthusiasm for the joint attack; these tactics are generally successful in creating more symmetric emotional engagement. Interestingly, in contrast to human children, chimpanzees fail to re-engage (human) partners in activities following interruptions (Warneken et al., 2006). Possibly, the finding that capuchins and humans, but not chimpanzees, exhibit partner re-engagement is evidence of convergent evolution between capuchins and humans regarding awareness of joint commitment towards common goals among partners. This would be consistent with evidence indicating convergent human-capuchin evolution regarding the importance of coalitionary aggression.

We observed considerable variation in (a) the ways various combinations of the basic behavioural elements described above were incorporated into a dyadic ritual, (b) the posture and gaze direction of the participants, (c) the extent to which dyads were temporally consistent in the form of their rituals, (d) the extent to which there was symmetric emotional engagement, and (e) the degree of turn-taking for those rituals that had multiple roles. However, structural commonalities in the rituals have led us to hypothesize that they share a common function (as bond-testing signals, Perry et al. 2003) and/or ontogenetic process. Capuchin monkeys normally behave at a rapid pace, both in their destructive foraging style and in their social interactions (e.g. rapid-fire grooming exchanges). Even while resting, their visual attention typically wanders, seeking new foraging opportunities or monitoring others’ social interactions. In striking contrast, their more creative social rituals proceed via slow, deliberate movements, and the participants’ faces bear almost trance-like expressions. Although participants rarely make eye-to-eye visual contact, one or both monkeys focuses visual attention on some body part of its partner, often for several minutes at a time. Sometimes both participants focus their attention jointly on an object. The amount of time and the sustained focus devoted to these rituals suggests that the two ritual partners value one another highly. Another common feature of these interactions is that they typically involve some risk or discomfort, e.g. a finger in someone’s mouth where it is at risk of being injured by teeth, or a finger in another monkey’s eye socket, so that a quick movement could scratch the cornea. One monkey often twists another’s body into positions that look distinctly uncomfortable. The monkeys’ enthusiasm for these uncomfortable and/or risky interactions is consistent with Zahavi’s “testing of a bond” theory (Zahavi, 1977). Behaviours that are risky, uncomfortable or disgusting will seem aversive when received from a non-favoured partner, but pleasurable when received from a favoured partner; the emotional response elicited by the bond-testing behaviour informs the tester about the state of the relationship. This theory (minus the emphasis on risk/discomfort as an adaptive design feature in the ritual) closely mirrors Collins’ ideas about interaction rituals, in which partners assess one another’s behavioural responses to their interactions with them, obtaining useful information about their relationship status and how the partner feels about them, relative to other partner options (Collins, 2004).

### 3.2 Social, ritual, and game network structure

In Figures 1a-c, circle nodes represent females and square nodes represent males that belong to one of the two matrilines (blue borders for the subordinate matriline and red for the dominant matriline) or have migrated into Flakes group (black border). In (a), nodes are coloured based on the results of the greedy community detection algorithm of the igraph package. This algorithm is trying to find dense subgraphs by optimizing the networks modularity score. Edge weights correspond to the proximity index of each dyad. In (b) and (c), edge weights represent the relative frequency with which monkeys engage in all rituals (b) and games (c).

Visual inspection of the proximity network (Fig. 1a) highlights the fact that matriline members tend to cluster together, with immigrant males being less social (aside from the current alpha male, who was HE till the end of 2007, QJ from the end of 2007 till mid-2016, and finally MX). However, the diagrams showing frequency of ritual participation show that some of the most isolated immigrant males (e.g. NP) are active ritual participants, particularly with regard to the games. Fig. 1d also highlights the lack of correspondence between social centrality and ritual participation.

### 3.3 Who performs these rituals, what are the performing dyads’ characteristics, and what does this tell us about ultimate function?

Table 2 presents the results of the three MLPE models used to predict ritual rates; graphical representations of these data are found in Figure 2, along with other details of the analysis. The ICC and tau values indicate that individual idiosyncracy did not explain much of the variance in ritual rates. The proportion of time a dyad spends in proximity is not a strong predictor of ritual rate; this is inconsistent with Hypothesis 1 (“bond establishment and maintenance”), but not necessarily inconsistent with the “bond-testing” hypothesis (H2). Consistent with both hypotheses, those dyads with higher quality relationships were slightly more likely to perform rituals. However, consistent more with the bond-testing than the bond maintenance hypothesis, dyads were more likely to perform rituals if they were in the range of RQI = 0.7 − 0.9 (30% of 254 dyads) than in the highest RQ values (7.5% of 212 dyads for which RQI > 0.9); none of the 14 dyads with RQI < 0.3 performed a ritual. Finally, the model using coalition formation rate as a fixed effect demonstrates a positive (though modest) relationship between coalition formation rate and rate of ritual performance; this is consistent with both Hypotheses 1 and 2.

**Table 2:** Results of three separate Maximum-Likelihood Population Effect (MLPE) models predicting ritual rate.

|  | Model 1<br>Proximity | Model 2<br>RQ | Model 3<br>Coalitions |
| --- | --- | --- | --- |
| intercept | 0.019 | 0.008 | 0.019 |
| estimate | -0.0014 | 0.089 | 0.11 |
| SE | 0.035 | 0.040 | 0.04 |
| P | 0.967 | 0.027 | 0.004 |
| 95% CI | -0.07–0.07 | 0.01–0.17 | 0.04–0.19 |
| $\sigma^2$ | 0.90 | 0.89 | 0.89 |
| $\tau_{00}$ individual | 0.05 | 0.06 | 0.05 |
| ICC | 0.05 | 0.06 | 0.05 |
| N individual dyads | 45,762 | 45,693 | 42,676 |

**Figure 2:**
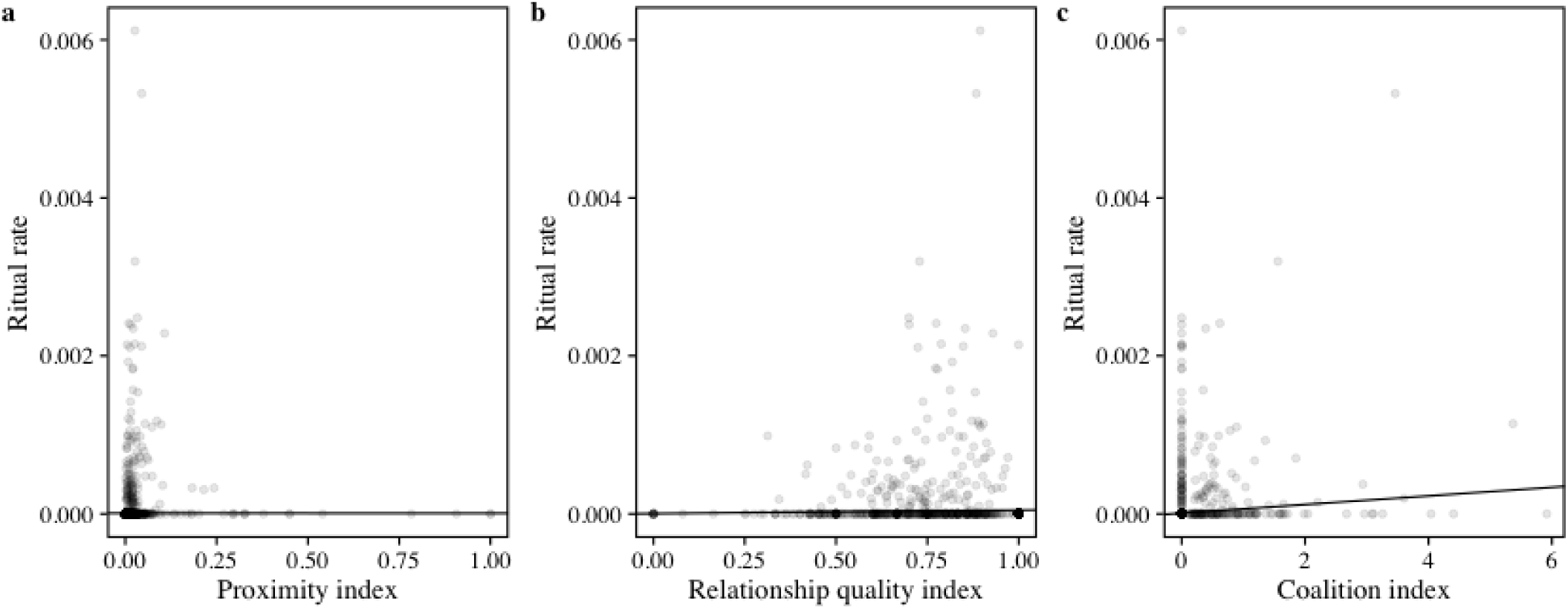
Graphical representations of the relationship between ritual performance rate (y-axis) and (a) physical proximity, (b) relationship quality, and (c) coalition index.

Examination of Figure 2a reveals that, consistent with the bond-testing hypothesis and inconsistent with the bonding hypothesis, most of the high values for rate of ritual performance are in the lower regions of the proximity index, i.e. in those pairs that spend little time together and hence may lack information about the current state of their relationship. In Figure 2b, the distribution of data points is fairly consistent with both hypotheses, but more so with the bond-testing hypothesis. Note that most of the high values for ritual performance are between 0.7 and 0.9 on the RQI scale, i.e. a zone in which dyads have pretty amiable relationships, but there are fewer high scores for ritual rate in the zone between 0.9 and 1.0 (highest quality relationships),despite the fact that ∼30% of dyads have RQI > 0.9.

In order to investigate when, developmentally, individuals began performing rituals, we examined a subset of the data: the ritual participation histories for the 17 monkeys (7 female, 10 male) who were immatures when first observed to participate in the toy and/or hair games, i.e. the rituals exhibiting the most complexity and requiring the most active participation. We found that individual capuchins were first seen to participate in some sort of ritual (game or non-game) at a mean age of 1.9 years (range 0.1-4.8 years), but were first seen to be active participants in games at a mean age of 3.2 years (0.7-7.3 years), with one female never becoming an active participant. The absence of these behaviours in younger individuals suggests that learning is involved in their production. For the 7 females, the “tutor” (i.e. the older monkey who was the other participant) was her (probable) father in 5 cases, an unrelated adult male in one case, and a female relative in the other case. Males had a wider variety of “tutors”, all of them male, including 2 fathers, 3 unrelated adult males, and in some cases other juvenile males very close to them in age.

## 4 DISCUSSION

In the previous sections we have described some unusual behavioural sequences observed in white-faced capuchin monkeys and provided an argument for calling these “rituals.” Our analysis supports the notion that these behaviours are relevant for dyadic bond-testing. In the following section we compare the observed rituals with human rituals and analyse form and function.

### 4.1 Comparisons of form and proximate causes

The form of the capuchin rituals described here bears some resemblance to other nonhuman primate rituals (e.g. baboon greetings) and to many types of human interaction rituals. As far as we can tell in the absence of similar methodologies across studies, it seems that the degree of behavioural variability in capuchin rituals is somewhat greater (i.e. less rigid and rule-bound) than in human rituals. The exaggeration of movement so typical of more species-stereotypical mammalian rituals (e.g. displays) is absent in capuchins. There is less obvious “framing” of the onset of rituals in capuchins than in humans, or even in baboons (Watanabe and Smuts, 1999). Though most of the dyadic rituals described in this paper start in the context of grooming, resting in contact, or slow motion play, there is no one behavioural or contextual element that reliably signals that a ritual is beginning or ending, even within a single dyad. It seems likely that the proximate trigger for these rituals is the monkeys’ perceived need for information about the status of the relationship, but we do not currently have a means of testing that hypothesis.

A commonality between capuchin rituals and human rituals is the attentional focus, which is often focused on a “sacred object,” i.e. an object that gains its value from the emotional charge acquired via its use in the ritual, rather than from any intrinsic utilitarian value (Collins, 2004; Goffman, 1967; Sosis and Alcorta, 2003). An important difference, however, is that the symbolic meaning of sacred objects in human rituals continues outside the context of the ritual; as far as we can tell, this is not true in capuchins.

### 4.2 Function, and the relationship between form and function

Whereas some theories regarding the evolution of ritual have focused more on the benefits relating to working memory (Rossano, 2011), others (e.g. Collins 2004; Zahavi 1977) have focused more on how the quality of the attention itself can serve as a signal of the partner’s current emotional and motivational state, which is relevant to assessing commitment to their relationship. It has been hypothesized, both for many types of human rituals (e.g. religious rituals: Bell 1997; Durkheim 1912; Rossano 2010; Sosis and Alcorta 2003) and for some types of nonhuman rituals (e.g. De Marco et al. 2014; Perry et al. 2003; Watanabe and Smuts 1999; Whitham and Maestripieri 2003), that ritual serves a social bonding function, enhancing feelings of solidarity, trust and desire to collaborate, or at least testing commitment to a particular group or partner(s). Although testing these functional hypotheses is difficult in both humans and nonhumans, the evidence from capuchins is generally consistent with the idea that capuchin dyadic rituals serve a bond-testing function. An important difference is that whereas most human rituals seem designed to promote group-wide solidarity, capuchin (and other nonhuman primate) rituals seem to operate at the dyadic level (Sosis and Alcorta, 2003), which has important implications for the relationship between form and function of rituals. Although capuchins do seem to have a strong sense of group identity (as exhibited by their xenophobia and collaborative aggression towards members of neighbouring capuchins groups, (Perry and Manson, 2008)), we have seen no examples of capuchin rituals in which all group members perform actions in strict unison, and capuchins very rarely cooperate as an entire group. Current theorizing about the function of ritual in humans also emphasizes the value of ritual for promoting adherence to group-specific social norms; possibly, capuchins lack such social norms.

The degree of rigidity in the form and ordering of the ritual actions is often considered a necessary diagnostic feature for rituals (Bell, 1997), and examination of this feature might provide insights into function. When a ritual’s function is group-wide bonding/identification, promoting group-wide cooperation, we should expect group-wide uniformity in the performance of a ritual. Our data make us sceptical that this is the function of capuchin rituals. In the capuchin data set, there was considerable inter-individual and between-dyad variation in the behavioural elements included in the ritual repertoire, and there was between-dyad variation in the level of mutual engagement and role reversals as well; section A2 of the Appendix describes some of this variation, discussing case studies of the ritual networks for four individuals. Capuchin rituals are more likely to be designed by natural and cultural selection to test and/or strengthen dyadic bonds, enabling individual monkeys and dyads to understand where they stand with regard to commitment and cooperation compared with other individuals and dyads within their social group. If this is correct, then we should expect to see high within-dyad uniformity, but less between-dyad uniformity than is seen in human rituals that are performed in groups. Indeed, following the logic of (Perry et al., 2003), between-dyad variation in the form of a ritual may be a design feature. The time required to devise a unique dyadic ritual would be non-transferable to other dyads, creating an opportunity cost that serves as an honest signal of commitment to that particular dyadic partner. Reminders of unique dyad-specific games played exclusively with a particular partner might create links between the past, present, and future of that dyadic relationship (a phenomenon akin to “traditionalism”, Bell 1997), in which the dyad-specific ritual behaviour may help create a mental representation of the social relationship. If this functional hypothesis is correct, then we would expect the following pattern of variation in capuchins: (a) increasing homogeneity within each dyadic ritual, as the partners come to an agreement of what roles and behavioural sequences characterize their unique dyadic ritual, (b) greater within-dyad homogeneity in form than between-dyadic homogeneity in form, even for dyads including one of the same individuals, (c) absence of within-group homogeneity in form, aside from the trivial similarities that come from the fact that independent inventions of rituals are constrained by the types of building blocks existing in the “Zone of Latent Solutions” (Tennie et al., 2009) for the species, and the obvious advantages of including behavioural elements that have Zahavian bond-testing qualities (i.e. are risky or uncomfortable for dyads with poor quality relationships, Zahavi 1977). Unfortunately our data set currently includes insufficient numbers of rituals for most dyads to test these hypotheses.

### 4.3 Ontogenetic and phylogenetic aspects

In both human religious rituals and nonhuman rituals, elements of feeding, drinking, and washing behaviours are often introduced into new contexts, i.e. taken from their original functional context and repurposed for communicative means. In the case of capuchin rituals, most of the behaviours come from the behavioural domains of (a) grooming, (b) extractive foraging (prying open or probing into holes and crevices with fingers, substituting a social partner’s body parts for the plant parts upon which these actions are performed in a foraging context), or (c) food sharing/tolerated theft (in which an individual exhibits close-range inspection of another’s hands or mouth and gently attempts to remove a piece of the food from the monkey in possession of the food; in the ritual case, a non-food item is substituted for food). When these behavioural elements are applied in the context of ritual, the presumed original functions of these actions (e.g. hygiene, in the case of grooming; nutritional gain, in the case of extractive foraging and tolerated food theft) are replaced by a new function, presumably related to the establishment, maintenance and/or testing of social bonds. It is not entirely clear on what time scale (ontogenetic or phylogenetic) this repurposing occurs in capuchins. Because not all individuals express the same rituals, and these rituals are generally developed and expressed later in life, it is likely that most individuals independently invent these rituals by repurposing behavioural elements and subsequently socially transmit them to partners via ontogenetic ritualization, i.e. that the borrowing occurs within the lifetime of an individual. Given the similarities in form across so many individuals and dyads, it seems likely that capuchins as a species (or genus) have evolved a proclivity to prefer to borrow these particular kinds of behaviours (i.e. elements from the grooming, extractive foraging and food transfer repertoires) rather than other behaviour types, due in part to the fact that they are descended from a long evolutionary line of animals that relies on extractive foraging and social learning about food that occurs in a scrounging context; this would be more of a phylogenetic argument.

### 4.4 Comparisons with human playground rituals

Besides these characteristics that are commonly aspects of definitions of ritual, the behaviours we describe here have additional characteristics that are part of Burghardt’s (2017) definition of play: (1) They appear to be spontaneous, pleasurable, rewarding and voluntary for at least one, and almost always both, members of the dyad performing them. (2) They are performed in the absence of any obvious acute or chronic stress, when the participants seem relatively relaxed. (3) Elements are often repeated within a single ritual performance or in subsequent performances by the same dyad, but not typically in rigid rhythmic or stereotypic form. Lack of immediate purpose is also a feature of definitions of both play and ritual (Burghardt, 2017). In some ways, capuchin rituals resemble human children’s playground rituals. Merker (2009) points out that whereas the motor details of children’s’ rituals are mainly arbitrary with respect to function, there is social pressure to do things in a particular way, and the propensity to care about these details, i.e. to conform, has a bond-testing function. That is, the obligatory stereotypy of the rituals makes it obvious when mistakes (deviations) occur, and to avoid making such mistakes, it is necessary to invest much time in practice. Learning the details of a dyadic or group greeting ritual, for instance, requires that the individual pay close attention over long periods of time and practice; this is a costly way of indicating investment in the relationship(s). Capuchin rituals are simpler than children’s hand-clapping games or secret handshakes, but they too seem to require extensive practice at mastering arbitrary details. The patterning of behavioural variation suggests that participants recall their usual roles with particular partners and repeat them, as if reaffirming their roles in this particular relationship.

## 5 CONCLUSIONS

The capuchin dyadic interaction rituals described here are characterized by a strong attentional focus on the partner’s body and/or a “sacred object”, repurposing of behavioural elements from the extractive foraging repertoire, and incorporation of risky or uncomfortable behaviours. The form of these behaviours makes them ideal as Zahavian bond testing rituals, but is also consistent with a bond maintenance hypothesis. The patterning of which dyads performs these rituals most often best supports the bond-testing hypothesis. The group solidarity hypothesis is supported neither by the form of the rituals (which are highly variable between dyads within the same group), nor by the temporal aspects, as these rituals are performed by dyads in isolation, rather than by many monkeys simultaneously. Although there is a fairly high degree of consistency within dyad regarding the behaviours performed, there is more creativity, less rhythm, and less precise replication of behavioural elements than is consistent with many definitions of ritual in the ethology, psychology and anthropology literatures.

## ACKNOWLEDGEMENT

We thank the following people for assistance with data collection, data cleaning and/or logistics: C. Angyal, A. Autor, B. Barrett, L. Beaudrot, M. Bergstrom, R. Berl, A. Bjorkman, L. Blankenship, T. Borcuch, J. Broesch, D. Bush, J. Butler, C. Carlson, S. Caro, A. Cobden, M. Corrales, J. Damm, A. Davis, C. deRango, C. Dillis, N. Donati, R. Dower, A. Duchesneau, K. Feilen, T. Fuentes Anaya, A. Fuentes, M. Fuentes, C. Gault, H. Gilkenson, I. Godoy, I. Gottlieb, J. Griciute, L.M. Guevara R., L. Hack, R. Hammond, S. Herbert, C. Hirsch, M. Hoffman, A. Hofner, C. Holman, S. Hyde, O. Jacobson, E. Johnson, K. Kajokaite, M. Kay, E. Kennedy, S. Kessler, W. Lammers, S. Leinwand, L. Johnson, D. Kerhoas-Essens, S. Koot, W. Krimmel, S. Lee, S. Lopez, S. MacCarter, J. Mackenzie, M. Mayer, F. McKibben, C. Mitchell, W. Meno, A. Mensing, M. Milstein, Y. Namba, A. Neyer, C. O’Connell, J.C. Ordoñez J., R. Popa, K. Ratliff, N. Roberts Buceta, E. Rothwell, J. Rottman, H. Ruffler, S. Sanford, C. M. Saul, A. Scott, I. Schamberg, E. Seabright, C. Schmitt, S. Schulze, J. Shih, S. Sita, W.Tucker, K. van Atta, J. Vandermeer, L. van Zuidam, J. Verge, A. Walker Bolton, M. White, E. Williams, L. Wolf, D. Wood, M. Ziegler. We are particularly grateful to W. Lammers and H. Gilkenson, for long-term management of the field site from 2003-2013. D. Cohen assisted with management of the database. Thanks to Colin Twomey, Andy Lin, and Bill Peterman for their feedback and help with the statistical analysis, and to J. Manson and 2 anonymous reviewers for comments. Permission to conduct the research was provided by the Costa Rican Park Service (SINAC and ACAT), and two private ranches (El Pelón de la Bajura and Ponderosa).

## SUPPLEMENTAL MATERIAL

All simulation code and data used in this paper will be stored on Dryad upon acceptance. Video files with ritual performances can be found online at https://lbmp.anthro.ucla.edu/research/social-traditions/.

## APPENDIX

### A1 TRANSCRIPTION OF VIDEO CLIPS DEMONSTRATING RITUALS

#### A1.1 LBMP - Capuchin Monkey Rituals - Clip1

The following example (and accompanying video) depict a ritual often performed by two immigrant males (BL and QJ), in which they take turns passing a “toy” (in this case a piece of bark, i.e. something that is not food) back and forth, each earnestly attempting to retrieve the “sacred object” from the other’s hand or mouth. Because they were born outside the study area, we do not know their exact ages, but they appear to be the same age (within a year of one another); BL joined Flakes in September 2004, group 5.5 months after QJ joined. They started devising rituals together in May 2005, initially giving one another “dental exams” which involved inserting their fingers in one another’s mouths. In February 2006 they started using one another’s hair as an object to pass back and forth, and by March 2006 they were also using “toys” such as the bark used in this example. They were mutually enthusiastic practitioners of these rituals until BL emigrated in November 2009. At the time of this particular interaction, QJ is the 2nd ranked male and BL the 3rd-ranked male in the group.

##### April 22, 2004, 9:10AM

At the start of the video, BL has his finger in QJ’s mouth (QJ on the right, BL on the left). BL uses his mouth and hand to try to pry QJ’s mouth open. QJ has a piece of bark in his mouth. [It isn’t obvious that this is bark until later in the clip.] QJ grabs BL’s hand (the one in his mouth) and removes BL’s hand from QJ’s mouth. QJ takes the piece of bark out of his mouth to show BL, perhaps to encourage BL to play the game. QJ is still holding BL’s hand. QJ puts the bark back in QJ’s mouth (or tries), but BL grabs QJ’s hand and tries to open it to extract what is there. They let go of one another’s hands, and then BL grabs QJ’s face, one hand braced against QJ forehead and the other prying QJ’s mouth open. BL uses his hand and mouth to pry open QJ’s mouth, using enough force to make QJ’s body sway. QJ adjusts BL’s hand in QJ’s mouth. Now BL uses both hands to pry QJ’s mouth open. BL gets the bark (or part of it) out of QJ’s mouth and puts it in BL’s mouth. Now each of them is using a hand to try to get bark from the other’s mouth. At 9:12:23, BL finally removes his hands, and QJ uses both hands to open BL’s mouth. BL pries QJ’s hands from BL’s face. They hold hands, and BL tries to extract the bark from QJ’s hand. QJ tries to get things out of BL’s hands. QJ succeeds in getting the bark from BL’s hand and puts it back in QJ’s mouth. BL is still working hard to get something out of QJ’s other hand. BL gets a piece of QJ’s bark and puts it in BL’s mouth. QJ grabs BL’s other hand and puts the bark in QJ’s mouth. QJ grabs something from BL’s lips and brings it to QJ’s mouth. BL lets some more bark protrude from BL’s mouth to show QJ, and QJ grabs that too, putting it in QJ’s mouth. BL grabs QJ’s head, turning it to face BL, and tries to pull the same bark from QJ’s mouth, using enough force to twist QJ’s body around a bit. BL’s finger seems to be clamped in QJ’s mouth. QJ eyes are closing. BL uses both his mouth and hands to try to open QJ’s mouth; QJ’s eyes are still closed. QJ grabs BL’s hand and pulls it out of QJ’s mouth. QJ grabs BL’s face and tries to open BL’s mouth. QJ seems to be chewing something. QJ’s finger is lodged in BL’s mouth. QJ tries to pull something from BL’s mouth, using both hands. When they are trying to get bark out of the other’s mouth, their visual attention is focused on the mouth rather than the eyes of the partner, throughout this interaction. QJ finally gets the bark out of BL’s mouth, puts it in QJ’s mouth, and chews it. BL immediately tries to retrieve it from QJ’s mouth. BL removes his hands and lies on his side, presenting for grooming while facing QJ. QJ tries to open BL’s mouth with both hands. BL holds QJ’s hands. BL inserts his hand in QJ’s mouth, then turns his attention to QJ’s hands, trying to pry one open. QJ tries to open BL’s hands. Their foreheads touch as they both focus on watching one another’s hand manipulations. QJ gets something out of BL’s hands, pops it in his (QJ’s) mouth, and chews it. BL immediately tries to get it out of QJ’s mouth. BL uses both hands to try to open QJ’s mouth, using much force. Finally, BL removes the bark and puts it in BL’s mouth. QJ scratches, then tries to open BL’s mouth. QJ removes his hand and scratches again. QJ sticks his finger in BL’s mouth. QJ uses both hands to pry open BL’s mouth. He fails, and BL uses both hands to try to open QJ’s mouth. BL gets something out of QJ’s mouth and chews it but continues to try to open QJ’s mouth. BL scratches his own head and drops his hands. QJ tries to pry open BL’s hand. BL’s attention strays, and he looks off in the distance. QJ jerks BL’s hand, as if to demand his attention. QJ lies down as if presenting for grooming but keeps his grip on BL’s hand. BL puts something in BL’s mouth. QJ tries to remove the bark from BL’s mouth with one hand, but while still reclining. Then QJ uses both hands. QJ stretches, inviting grooming. BL grooms QJ’s chest. BL stops grooming and turns away from QJ. BL is still playing with the bark. QJ grabs BL’s tail and tugs it 3 times, but BL doesn’t turn around. BL self-grooms and turns back to face QJ. BL flops down, presenting for grooming to QJ. It is not clear who ends the interaction sequence, because the video clip ends here, at 9:22.

#### A1.2 LBMP - Capuchin Monkey Rituals - Clip2

##### March 13, 2012

In this clip we see Minstrel (‘MI’, the alpha female) and her maternal half-sister Mead (‘ME’, who is two years younger), performing a ritual that is highly typical of their relationship (see section A2.1 of the Appendix for more details regarding their relationship). Minstrel, on the left, is clutching Mead’s hand and inserting it in her (Minstrel’s) mouth. Mead reclines, passively accepting this. Minstrel readjusts Mead’s finger, inserting it deeper into her mouth on the other side, and then switches back to the original position. This is just one of multiple clips from this ritual, but it is fairly typical both of this bout and of the relationship more generally.

### A2 BETWEEN-DYAD VARIATION IN RITUAL PERFORMANCE

#### A2.1 Case studies

As can be seen by visually inspecting the tab called “ritual details” in the RawData.xlsx file of the ESM, there was considerable variation between dyads regarding the elements incorporated into their rituals, and also variation in the degree of mutual enthusiasm/engagement. Limited space precludes presentation of the data for all dyads, but here we present some illustrative cases from individuals who contributed large sample sizes to the data set, focusing on the behaviour of four individuals: two immigrant males (QJ and NP) and two females (sisters ME & MI).

NP, a highly peripheral and shy male, contributed 57 rituals to the data set (2 of them failed attempts to engage others), and participated in rituals with 15 different monkeys, though he focused primarily on three of the older juvenile males. Of his 57 rituals, 5 involved insertion of fingers in or on the nose, 20 involved insertion of fingers in mouths, 3 were “dental exams”, 3 involved eye-poking, 2 involved sucking a body part other than the finger, and the rest were games in which an object (hair in 12 cases, a “toy” in 27 cases) were passed back and forth. Comparison of the network diagrams for physical proximity and ritual performance (Fig. 1) highlights how extensive NP’s ritual network was, despite his low rates of physical association with his group-mates.

QJ, an immigrant male who became the alpha male in 2007, was a far more socially central male than NP, and contributed 92 rituals to the data set. These were spread over 21 monkeys (11 females, 10 males), all but two of whom (the eldest females of each matriline) were younger than him. His most frequent partner by far was BL, an unrelated adult male who performed 21 rituals with him (one of them displayed in the video clip described in Appendix A1.1); all involved some sort of exploration of the partner’s mouth, and 9 involved a “toy” or hair game; in general, his rituals with BL were characterized by a high degree of turn-taking and mutual engagement. He also performed 5 or more rituals each with TU (an unrelated male), his son YJ, and his three daughters (BW, LD and IE). His relationship with his daughter BW was noteworthy for the active role she played in manipulating various parts of QJ’s head. In only one of these was an object involved (in which BW was trying to remove hair from QJ’s mouth).

The dyad consisting of Minstrel (MI, alpha female) and her half-sister Mead (ME, two years younger than Minstrel), contributed 72 rituals to the data set beginning in Feb 2004 when Mead was four years old and ending in January 2015. There was remarkable similarity in form across these events. Typically, Minstrel grabbed her sister’s hand and inserted Mead’s fingers into Minstrel’s nostrils and/or mouth. In all cases in which we were certain of the initiator, it was Minstrel; Minstrel terminated the interaction in 57% of the 30 instances for which we felt comfortable designating a terminator. Although Mead was an involved participant for the grooming portion of the ritual, she was far less engaged in the portion that involved insertion of her fingers in Minstrel’s orifices. Mead either lay there passively until Minstrel was done with her fingers, or, in some cases, actively resisted the interaction, struggling to adopt more comfortable positions than those favoured by Minstrel. In only three cases did Mead attempt to insert Minstrel’s finger in Mead’s mouth or nose. This pair was a sharp contrast to some of the male-male dyads (such as QJ-BL, described in the Appendix A1.1) that had more complex rituals characterized by high emotional engagement on both sides, and active turn-taking in which roles were reversed. This contrast is evident by comparing the two video clips described in the Appendix section A1.1 and A1.2.

Interestingly, although Mead seemed unenthusiastic about hand-sniffing and finger-sucking with Minstrel and almost exclusively had the role of “finger-donor”, she practiced these behaviours with several younger members of her family, and took the role of the more active participant (taking their fingers and inserting them into her mouth or nose, despite some resistance from them). Mead had 15 other relationships with one more ritual. In only three of these did she sometimes assume the role of the monkey whose finger was sucked or sniffed. In at least 8 of these relationships, it was clear that Mead was directing the action, and taking the hands of her partners to insert in her nose or mouth. Although it would be tempting, based on the patterning of Mead’s interactions alone, to say that dominance rank or age was the factor determining who sniffed or sucked on whose body parts, this pattern did not hold true for the group more generally. Intriguingly, HE, the male who was alpha male from 2004-2007 (and who remained in the group after being deposed) participated in a ritual only once, as a passive participant with his son, even though all of the other adult and adolescent males were enthusiastic practitioners.

### A3 ANALYSES OF PREDICTORS OF WHICH DYADS PERFORM RITUALS: ALTERNATIVE ANALYTIC APPROACHES

#### A3.1 Mixed effects negative binomial regression

Our first approach was to conduct mixed effects negative binomial regression models (in Stata 13.1), with crossed random effects for the two members of each dyad. The results were similar to those described in section A3.2, and also for the MLPE models, i.e. no significant effect of proximity, but statistically significant (albeit small) positive effects of relationship quality (RQI) and coalition rate on the rate of ritual performance. However, this approach does not take possible effects of the monkey identities into account.

#### A3.2 Negative binomial regression model

Our second approach, adapted from the econometrics literature, was to analyse the data using negative binomial regression models (in Stata 13.1). We used three different models for the three independent variables: proximity index (PI), relationship quality index (RQI), and coalition index. In each of these models we used a fixed effects intercept for each individual monkey, i.e. we created a dummy variable that represented whether a monkey was a member of a dyad. The response variable, in all cases, was the count of rituals performed by this dyad, with the exposure being the sum total of group scans and point samples collected on the two partners in the ritual. For the proximity and RQI models, we also ran versions in which each row was a dyad-year (i.e. having repeated measures for each dyad), because we were concerned that the quality of relationships might change over time and that this variation (and its relationship to ritual performance rate) would be lost by pooling large time periods. In these models that had repeated measures for dyads, we clustered robust SE on dyads.

The model for RQI (pooled over the entire observation time) had 693 observations (we dropped values with zero social interactions). There was a significant positive relationship between RQI and ritual rate (coeff. = 2.18, SE = 0.38, z = 5.8, P < 0.001, 95% CI 1.44 − 2.92). Results looked similar when each data point was a dyad-year (coeff. = 0.76, robust SE = 0.17, z = 4.36, P < 0.001, 95% CI 0.42 − 1.10).

The proximity model was more difficult to understand. When we pooled the data over the entire observation time, we found a significant positive relationship (coeff. = 7.37, SE = 2.11, z = 3.50, P < 0.001, 95% CI 3.24 − 11.50). However, this only occurred in very low values of the proximity index starting to level off or decline around the value of x = 0.02. Visual inspection of the graphical output indicated that even if there was a statistically significant relationship, it was not biologically significant. This result was an outlier in that other modelling approaches did not find a significant effect of proximity. In the model where time was pooled for each dyad and each year, the strength of this statistical association was much weaker (coeff. = 0.82, robust SE = 0.89, z = 0.92, P = 0.36, 95%CI −0.93 – 2.56).

For the coalition index model we found a positive correlation with ritual rate (coeff. = 0.37, SE = 0.16, z = 2.28, P = 0.02, 95% CI 0.053 − 0.696). There were several extreme values, for example, for dyads that performed many rituals but no coalitions or vice versa.

#### A3.3 Mantel test

However, neither of the two previous approaches preserved dyadic information. We used two further tests that preserve this information: Mantel test and MLPE (see the main text) models. The Mantel test compares two matrices. Our raw data are column entries of, e.g. ritual rates, for various dyads. We turned these values into matrices by creating an empty square matrix with row and column numbers that were equal to the number of unique identifiers in all dyads. We then filled in the column information in the correct cell for each dyad.

The Mantel test provides a Z-statistic, which is equal to the sum of the products of the corresponding elements of each matrix:

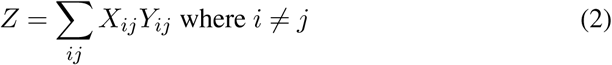

**Table S1:** Mantel test results. Results shown are for 5,000 permutations of the matrix pair in each row.

|  | Z | Coefficient | p |
| --- | --- | --- | --- |
| Ritual ~ Proximity | 0.003 | 0.006 | 0.227 |
| Ritual ~ RQI | 0.077 | 0.022 | <0.001 |
| Ritual ~ Coalition | 0.048 | 0.127 | <0.01 |

However, this value is highly dependent on the scale of the data. Therefore, we also calculated the Pearson correlation coefficient between the two matrices X and Y as:

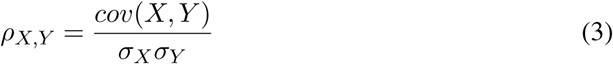

The Mantel test permutates one of the two matrices repeatedly. With each permutation, a correlation coefficient between the two matrices is calculated. This provides a distribution of correlation coefficients based on random permutations. The p value then represents the chance that the actual observed correlation coefficient between the two original matrices is based on chance. In Table S1 of the Appendix, we summarize the results of the Mantel test we ran. Again, the results show that proximity has no effect on ritual count, whereas relationship quality has a small but significant, and coalitions a larger and significant effect. However, because the Mantel test does not accept missing data, we had to create matrices that contained structural zeros for those dyads that were not co-resident. This is not ideal, which led us to the final set of models, the MLPE models described in the Results section of the main text, which do not require use of structural zeros.

The code for running these models in Stata and R is included in the supplementary materials.

